# Assessment of population dynamics of endangered *T. putitora* (Ham, 1822) in the breeding and nursery grounds of Western Himalaya

**DOI:** 10.1101/2023.07.06.547904

**Authors:** Priyanka Rana, Prakash Nautiyal

## Abstract

The present study analyzed the population characteristics of *Tor putitora* in their breeding and nursery grounds located in Western Himalayan region. From a sample size of 400 individuals, growth parameters such as asymptotic length, growth rate and growth performance index were estimated as 80.01 cm, 0.810 per year and 3.71 per year while the potential longevity was estimated as 4.6 years. The mortality parameters such as total, natural and fishing mortality were estimated as 2.88 /year, 1.05/ year and 1.83 / year respectively. The length at 25 %, 50% and 75% susceptibility to capture was estimated as 40.87 cm, 38.11 cm and 35.34 cm respectively. The fishing mortality and exploitation beyond optimum level and maximum sustainable exploitation ratio (Emax) indicate the overly exploitation of Mahseer resources in their breeding and nursery grounds. This indicates that the examined breeding and nursery grounds in Western Himalaya are facing intense “growth and recruitment overfishing” which may result in stock collapse in the forthcoming years.

## Introduction

The *Tor putitora* (Ham. 1822), commonly known as the Himalayan and Golden Mahseer, is a stenothermal and rheophilic species that primarily inhabits cold-water regions within the Himalayan Range. It is an endemic species to this region and has garnered considerable popularity due to its sporty behavior, which has been observed and admired by individuals around the world. It is also considered among one the largest freshwater cyprinids.

The Himalayan Mahseer undergoes a tri-phased migration as a means of sustaining their life-history characteristics, which include food resource management, overwintering/learning imprints, and spawning. (Nautiyal et al., 2001). This migratory phase is the major concern for the mahseer population as most of them get caught during their journey (Shreshta, 1997). Beside this, the other factors such as blockage of migratory path due to dam and reservoirs, regulated water flow, habitat destruction, pollution are the other major factor that are contributing toward population decline of Mahseer, which is already categorized as endangered species (Jha et al., 2018).

Therefore, in the context of current exploitation and man-made hinderance toward the mahseer population, it is imperative to implement a continuous assessment and monitoring program to ensure the sustainable management of Mahseer resources. In this regard, the study examines the fundamental population parameters along with current exploitation status of *T. putitora*. The study utilizes length frequency distribution (LFD), biometric index to analyze the various aspects of reproductive biology such as growth, mortality, asymptotic length (maximum length), potential longevity, length at first maturity and recruitment rate of Himalayan Mahseer. Reproductive biology provides valuable information about the reproductive processes, behaviors, and strategies of fish species to maintain the maximum sustainable yield of their wild stock (Ahmed et al., 2012). Furthermore, it has been posited that an accurate assessment of life-history traits can be estimated through an evaluation of early life stage and juveniles, as opposed to solely relying on adult-size groups (Cowan and Shaw, 2009). Hence, the present study is restricted to the breeding and nursery grounds, encompassing a variety of size compositions ranging from younger stages to brooders of Gangetic stock of Himalayan Mahseer. This study aims to provide the necessary information for the sustainable management of endangered Mahseer population in the breeding and nursery grounds of Western Himalaya.

## Study Area

The Nayar, located in the Western Himalayan region, is a tributary of the Ganga that is fed by springs and located approximately 10 kilometers downstream of Devprayag, functions as a breeding and nursery ground for *T. putitora* stock. The Nayar originates from the Dudhatoli peaks of Pauri Garhwal district in Uttarakhand, where it takes the form of Eastern and Western Nayar (as illustrated in Fig. 1). Ultimately, these two rivers converge at Batkul ka Sain, thereby forming the primary channel of the Nayar, which eventually merges with the Ganga at Vyasghat. The life cycle of the Himalayan Mahseer is intricately linked with the interdependent tributaries of the Ganga, namely Alaknanda, Bhagirathi, Song, and Nayar, as espoused by Nautiyal et al., (2001). However, it is only the Song and Nayar rivers that are situated close to the Gangetic Mahseer population, which serves as a breeding ground (Nautiyal, 2002) and nurtures juveniles as nurseries through the provision of a bountiful food supply (Nautiyal, 2013). As per the 2013 CPCB report, the Song River is currently encountering an alarming rate of sewage and pollutant load. In contrast, situated approximately 40 kilometers upstream of the foothill Ganga, the Nayar serves as a crucial breeding ground for the Mahseer population, benefiting from comparatively lesser human interventions.

**Figure 1.**
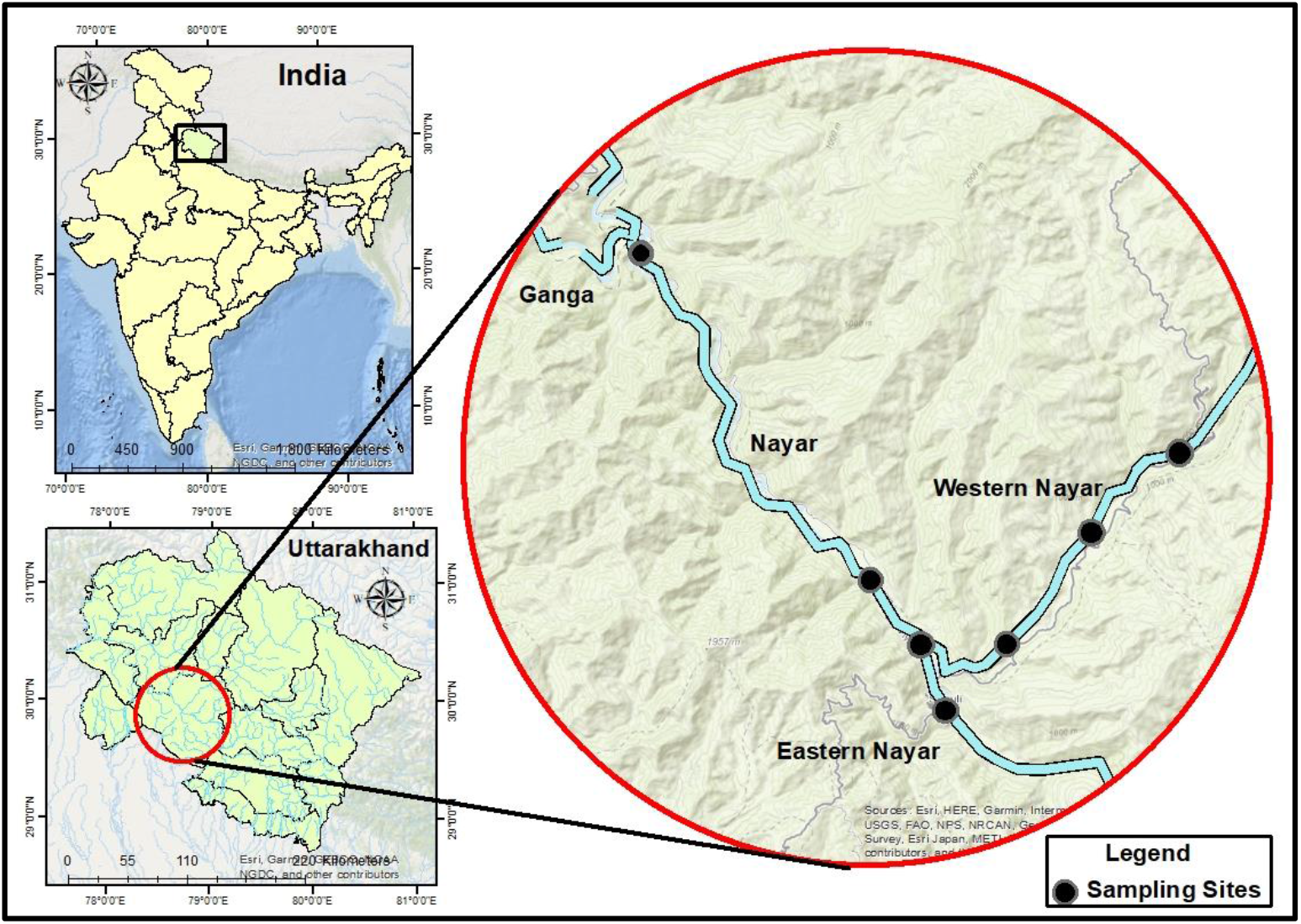
Study area and sampling locations.

## Method

The current investigation was implemented over a period of 4 months spanning from July 2022 to October 2022 which is regarded as the breeding period of Himalayan Mahseer. During this period, the different size-groups of Mahseer are easily available classifying from larval stages, adolescents and mature brooder stages. A careful selection of sampling sites within the Nayar was made, from where fish samples could be collected on a weekly basis via local fishermen (Fig I). All *T. putitora* specimens were thoroughly cleansed utilizing a cotton cloth. Upon retrieval, the total length was measured to the nearest 0.1 cm with the aid of measuring tape. The different size-classes examined during the study period were subsequently grouped into size classes of 10 cm interval on the basis of their length.

### Growth and mortality parameters

The growth and mortality parameters of *T. putitora* were examined by analyzing the length frequency data through diverse models using FiSAT II software (FAO-ICLARM stock assessment tool) version 1.2.2. Specifically, the von-Bertalanffy growth parameters (VBGF), comprising of asymptotic length (L∞) and growth rate (K), have been evaluated using ELEFAN II method. The asymptotic length (L∞) represents a theoretical or predicted value that is derived through the predicted growth rate and potential longevity (t_max_) of the fish within the stock. Theoretical age at zero length (t0) and growth performance index (φ) has been evaluated as per (Pauly, 1984);

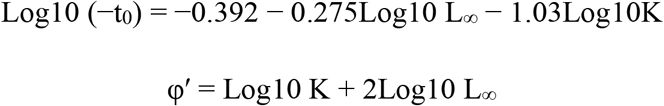

While the value of potential longevity (t_max_) has been calculated using,

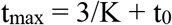

For the estimation of growth rate against the increasing mean length of Himalayan Mahseer, the Bhattacharya method given by FiSAT II software was employed and thereafter, function of linking means were performed.

Subsequently, the utilization of FiSAT II software facilitated the execution of the VBGF graph, recruitment patterns, and virtual population analysis. The length-converted catch curve model provided by FiSAT II software was utilized to compute total mortality (Z), natural mortality (M), and fishing mortality (F). The average annual temperature of the designated stretch was recorded at 21.9°C. Annual mortality (A) was calculated using the equation (Ricker, 1975);

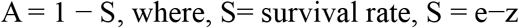

### Probability of capture

The probability of capture for fish susceptible to fishing gears (mostly cast nets) at different lengths was estimated at 25%, 50%, and 75% (i.e., L25, L50, L75). The ascending left component of the length-converted catch curve was used to assess the probability of capture of the fish (Gayanilo et al., 2005).

### Exploitation

The exploitation ratio (E) and rate of exploitation (U) of the given stock was calculated using the formula (Pauly, 1984);

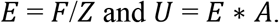

The exploitation ratio helps to assess whether the stock is under-exploited (E< 0.5) or over-exploited (E> 0.5) in the selected stretch (Gulland, 1971). The exploitation rate is considered optimum when fishing mortality is 40% of the natural mortality i.e., *Fopt* = 0.4 ∗ *M* (Pauly, 1984). Relative yield per recruit and relative biomass per recruit were also analyzed using the Knife edge selection of the Beverton and Holt model in the FiSAT II software.

### Recruitment pattern

The recruitment stock refers to either the number of fish that enter into the fishing stock or the size of the stock that is being utilized for fishing purposes (Gulland, 1965). The length at first recruitment (Lr) of the observed stock was calculated using the midpoint of the lowest class interval among observed length-frequency samples (Gheshalgi et al. 2012). The annual major and minor recruitment pulses along with their strength were calculated using FiSAT II software.

## Results

During the study period, about 400 specimens of *T. putitora* were examined. Notably, the examined size classes had the smallest mid-length of 6.2 cm. The dimensions of *T. putitora* specimens exhibited a range from 6.2 to 79.8 cm, comprising both resident stock, which measures less than 25 cm (present throughout the year), and seasonal stock of migratory adults and brooders, which measure greater than 25 cm. The majority of individuals constituted the size group ranging from 10-15 cm, succeeded by those in the size groups of 5-10 cm and 15-20 cm (as indicated in Figure 2). The size group exceeding 25 cm constituted a relatively lower proportion of the entire cohort since the large-sized brooders migrate for spawning during the initiation of the monsoon season, as stated by Nautiyal et al. (2001).

**Figure 2.**
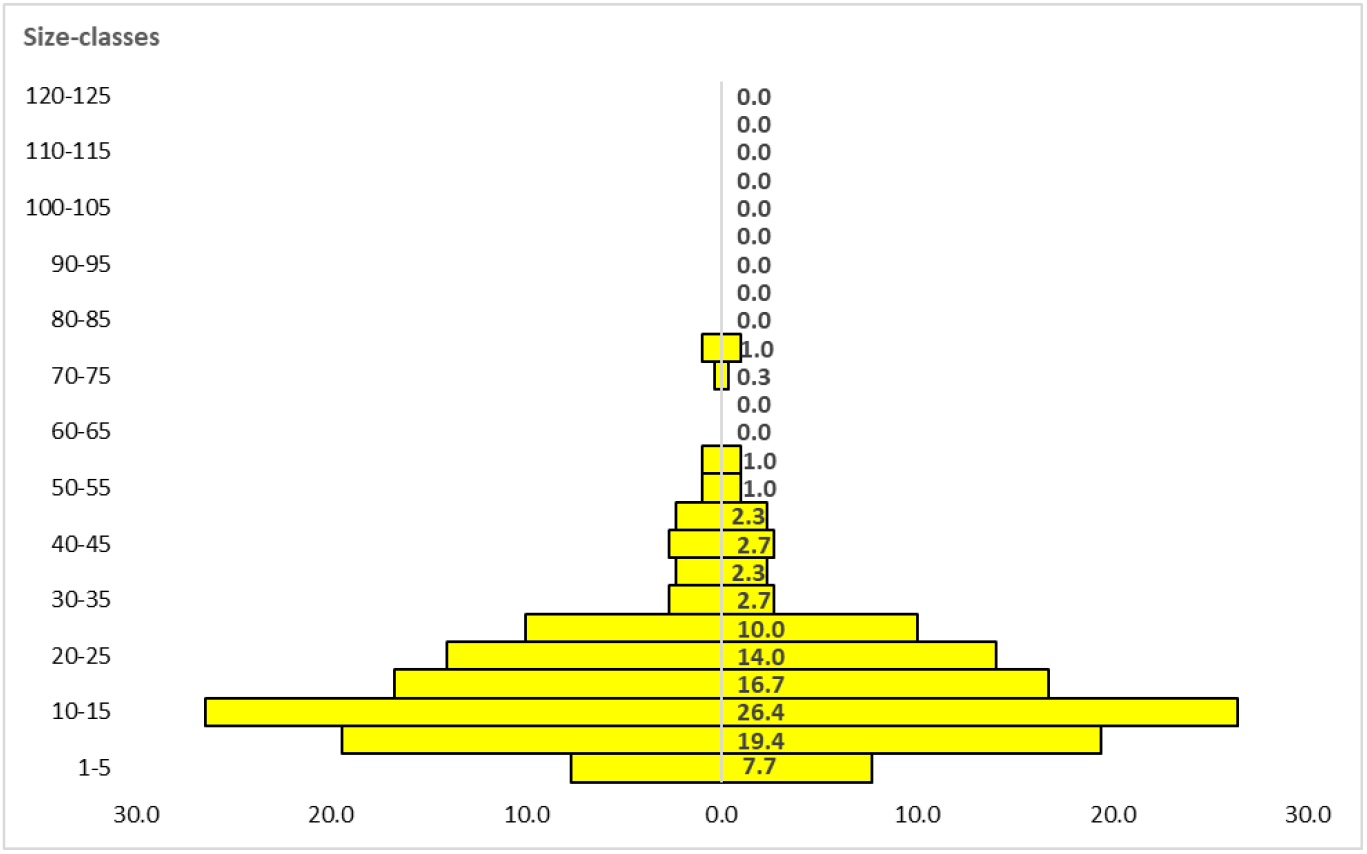
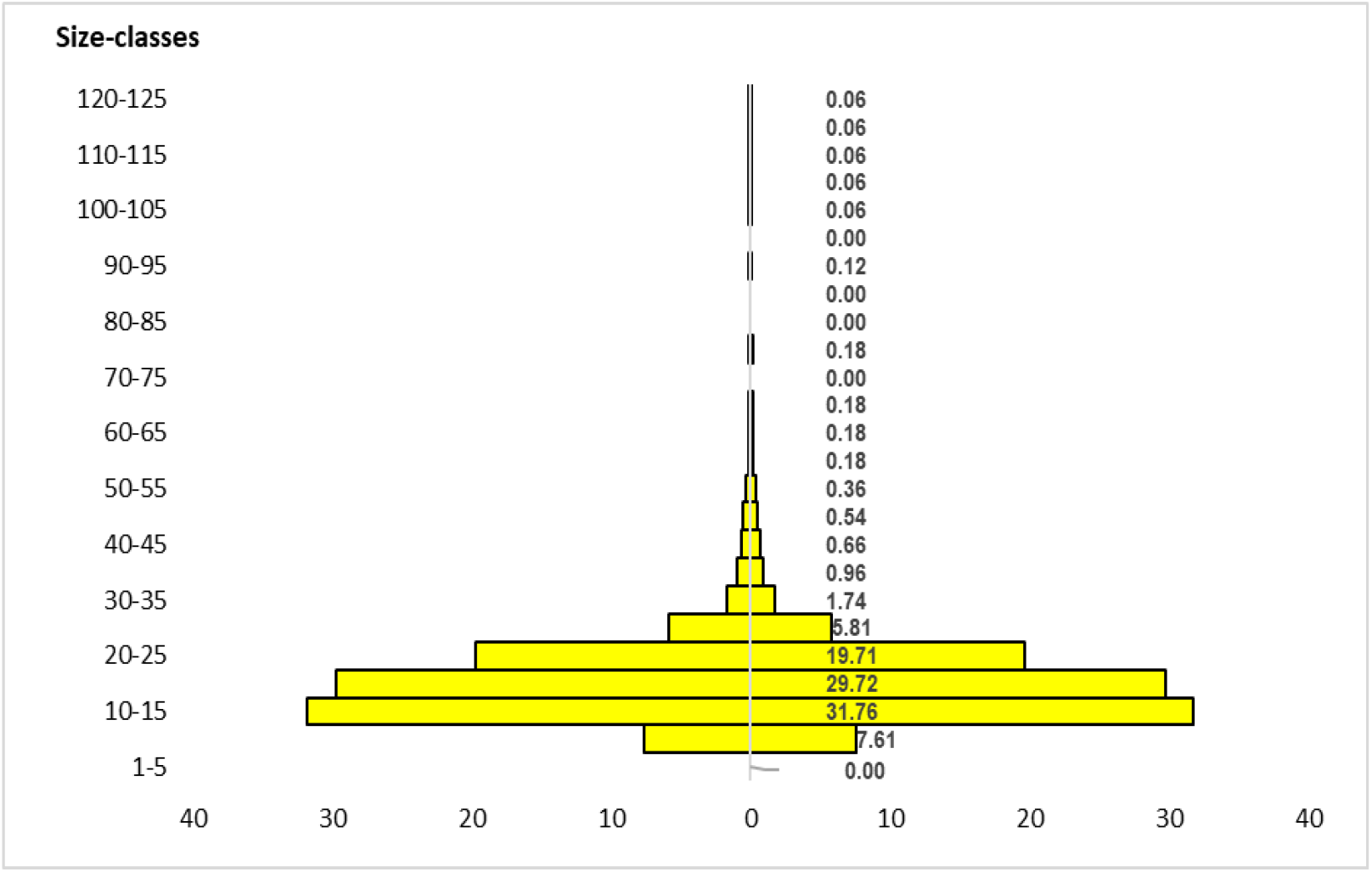
Size composition of *T. putitora* in the Nayar river.

### Growth parameters

The asymptotic length (L_∞_) was estimated as 80.01 cm with the growth rate (k) of 1.1 per year while growth performance index (Φ) was calculated as 3.710 per year. The estimated age of the hatchling (t_0_) was calculated as -0.82 per year while potential life-span (t_max_)was calculated as 4.6 years.

A slow growth rate against the increasing mean length of *T. putitora* has been recorded for the present cohort (Fig. 3). However, continuous growth was observed throughout the year as seasonal growth oscillations were found to be significant and strong (C < 1) indicating influence of seasonality on the growth of the species. The growth of the juveniles was seen continuous throughout the year while during May growth gets reduced but still continuous and reaches to its maximum in October (Fig. 4).

**Figure 3.**
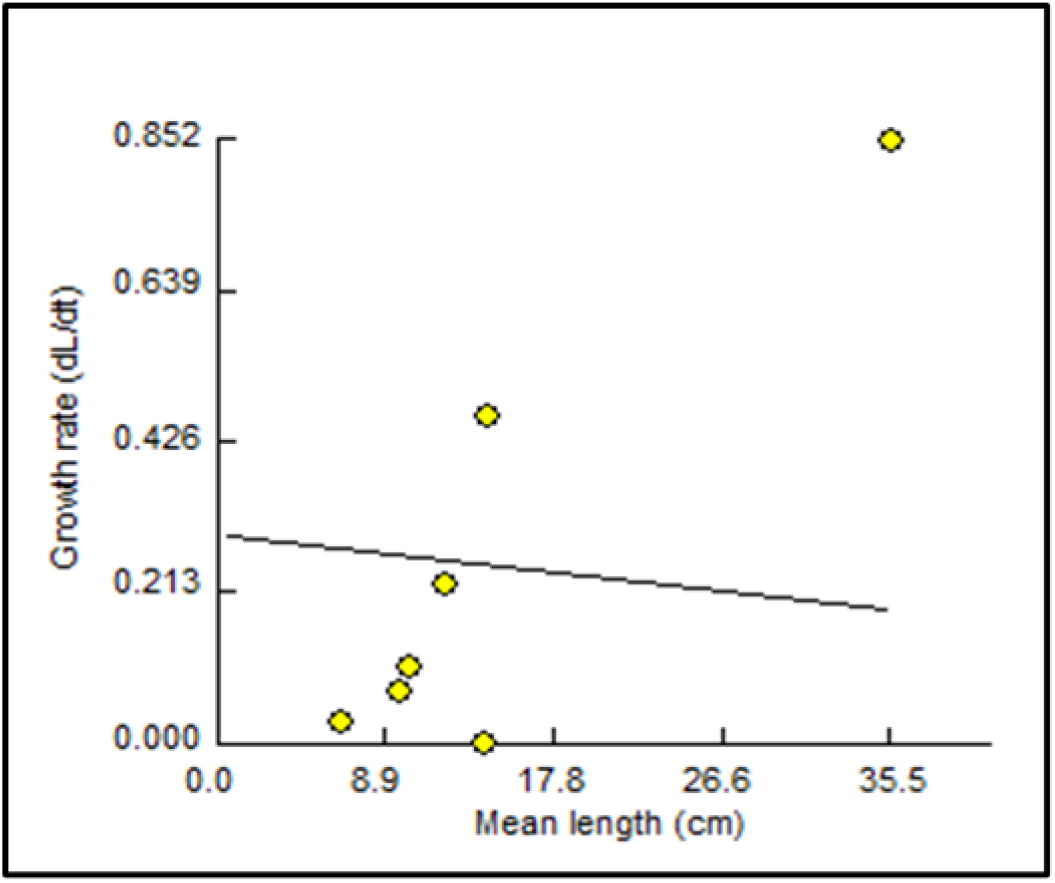
Gulland and Holland plot for growth rate determination against the mean length of *T. putitora*.

**Figure 4.**
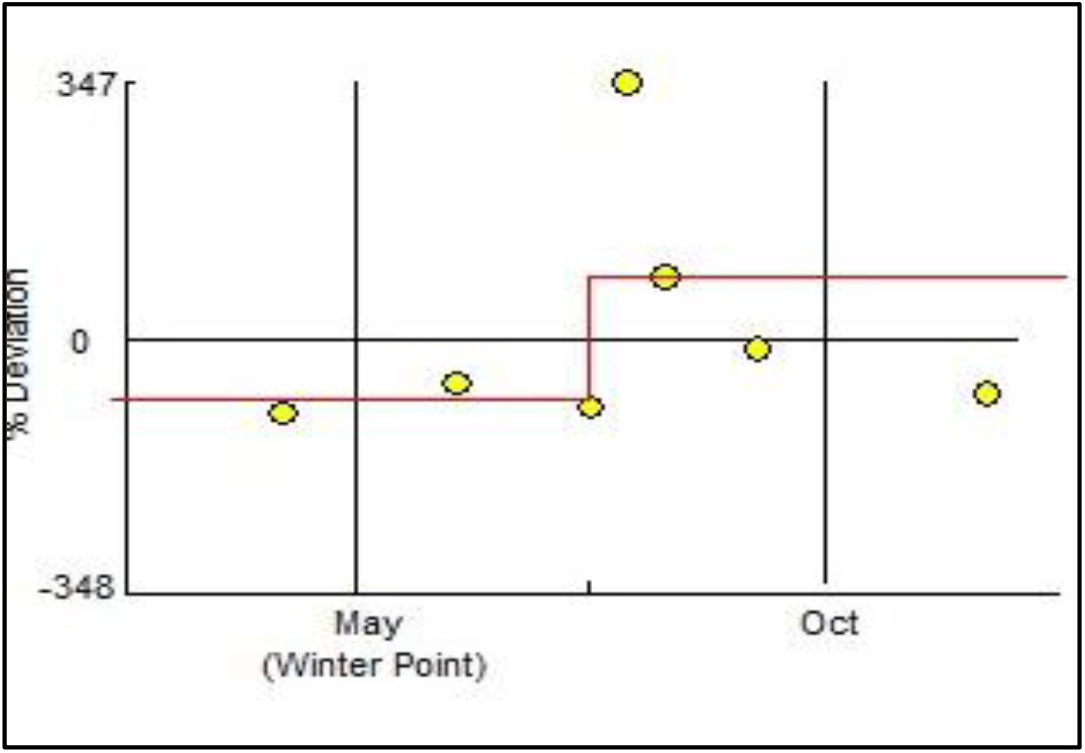
Seasonal growth oscillations of *T. putitora*.

### Mortality parameters

The total mortality (Z) was estimated as 2.88 where, natural mortality accounted about 1.05 and that for fishing mortality was 1.83 (Fig. 5). The survival rate (S) was estimated at 0.05 while that of annual mortality (A) was estimated as 0.94. Here, the optimum fishing mortality was estimated at 0.42 while exploitation ratio was estimated as 0.67. Therefore, the exploitation ratio and fishing mortality has surpassed the optimum level.

**Figure 5.**
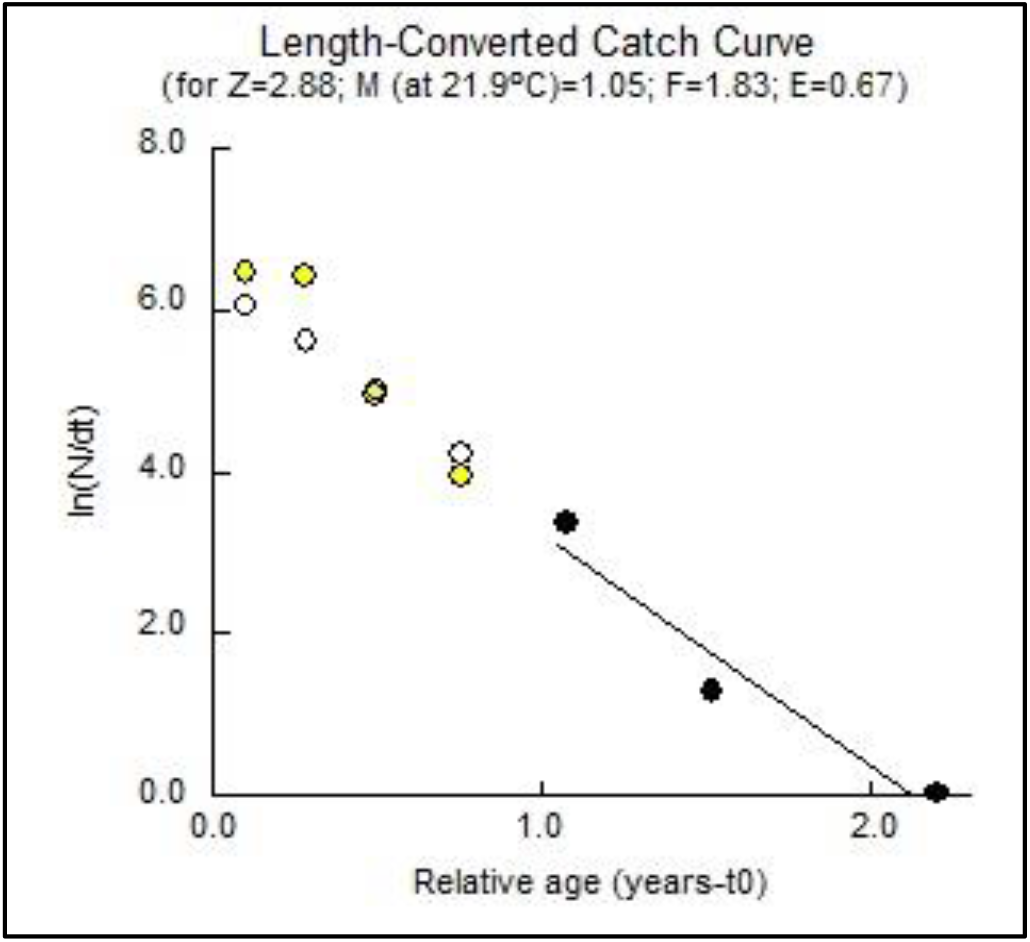
Mortality parameters of *T. putitora*.

### Probability of capture

The probability of capture extrapolated curve depicts that fishes measuring less than 16.20 cm are highly susceptible to being captured (as shown in Figure 6). However, in the current cohort, fishes measuring 10-15 cm constituted 30.98 5 % of the total catch, whereas those measuring 15-20 cm constituted 27. 98 %. Therefore, the fishes measuring below 20 cm formed about 65.4 % of the total catch.

**Figure 6.**
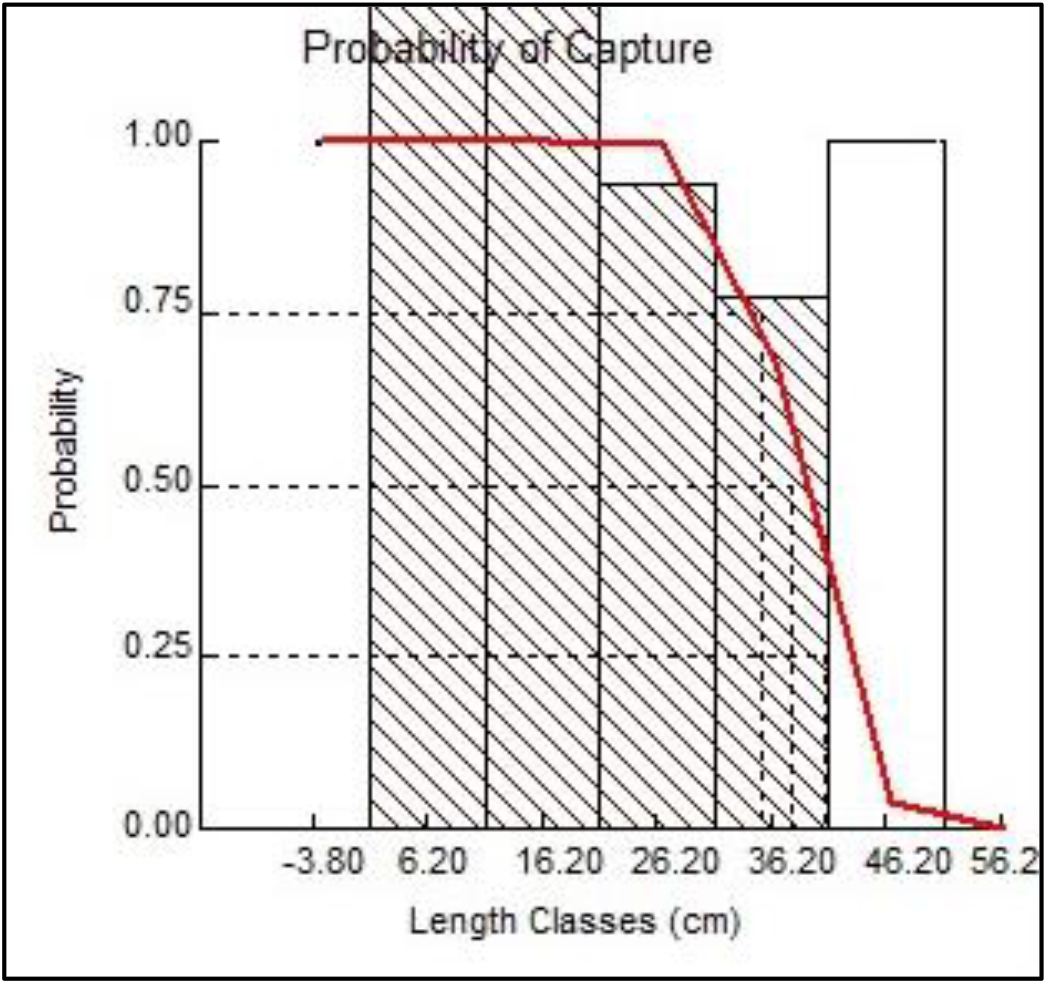
Probability of capture at different 25%, 50% and 75% susceptibility level.

The graph of probability of capture estimated L_25_, L_50_ and L_75_ as 40.87 cm, 38.11 cm and 35.34 cm respectively (Fig. 6). The virtual population analysis of have shown high fishing mortality of juveniles measuring between 16.2 to 26.1cm. However, another peak was observed in the size group measuring above 66.2 cm (Fig. 7).

**Figure 7.**
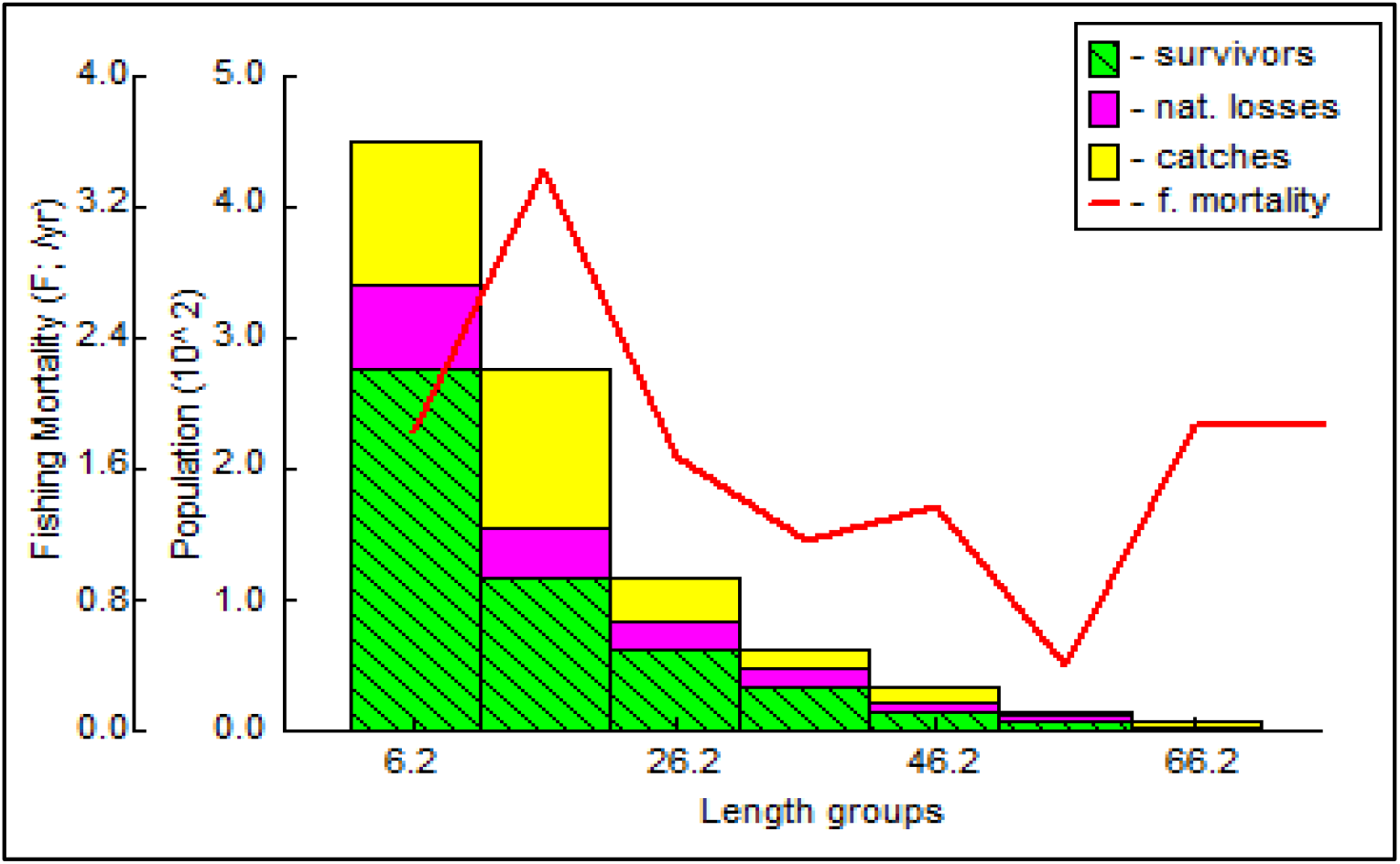
Virtual population analysis (VPA) of *T. putitora*.

## Exploitation

The graph of relative yield or biomass per recruit has estimated E_10_, E_50_ and E_max_. as 0.355, 0.278 and 0.421 respectively (Fig. 8). The predicted Emax therefore, found below the current exploitation ration which further indicate the overexploitation of *T. putitora* stock.

**Figure 8.**
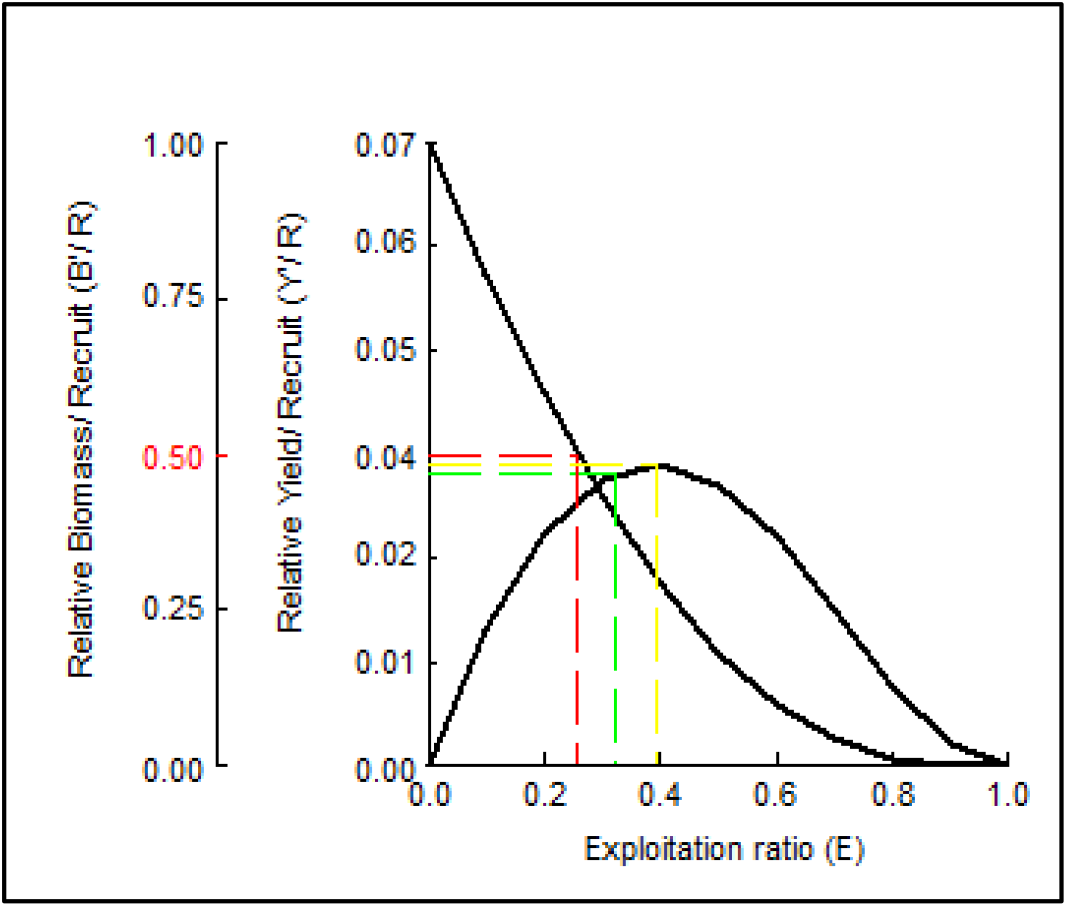
Relative yield or biomass per recruit analysis of *T. putitora*.

The relationship between the relative yield per recruit and biomass per recruit with exploitation ratio for both studied years is depicted by the Beverton and Holt plot. The input parameters, namely Lc / L∞ and M/K ratio, were estimated as 0.47, 1.2 respectively.

## Recruitment

The initial recruitment length Himalayan Mahseer was determined as 11.15 cm. This indicates an increase in the minimum length of the fish being exploited for fishing purposes. The recruitment process was observed to occur consistently throughout the year, demonstrating an efficient and effective recruitment cycle (Amponsah et al., 2016). The graph of recruitment has shown major recruitment pulses in January with 24.23% followed by February with 22.39 % (Fig. 9).

**Figure 9.**
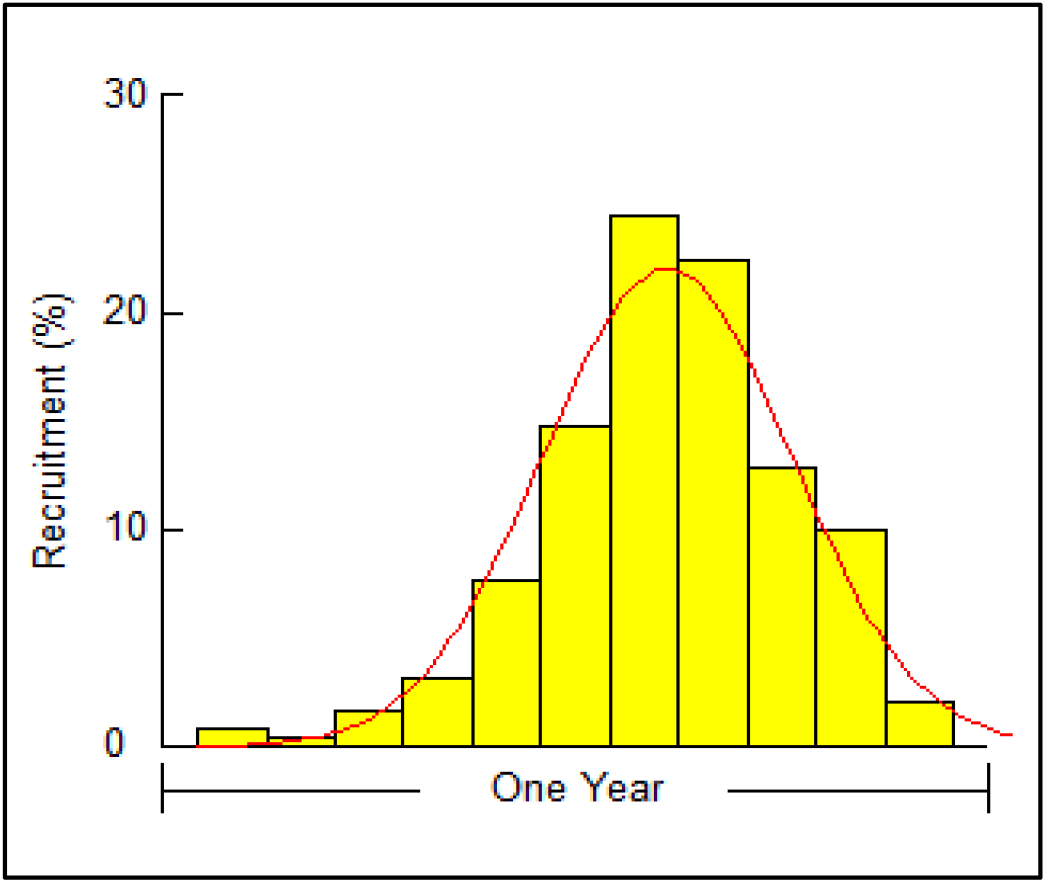
Recruitment pattern of *T. putitora* in the Nayar.

## Discussion

The present stock has revealed inverse relationship between asymptotic length and growth rate as also stated by Beverton and Holt (1957). This indicate that the species with relatively higher high asymptotic length usually possesses lower growth rate to attain their maximum length. This also denote their maximum life span compared to those with higher growth rate. For examples, species such as *T. tor*, Brown trout, Rainbow trout which grow more rapidly have comparatively shorter life span (Nautiyal et al., 2008). Earlier, the asymptotic length for this stock was estimated as 137.7 cm (Nautiyal et al., 2008) which has now been reduced to 80 cm.

Although in Gangetic stock, Mahseer with an age of 17^+^ years have been observed but the maximum lifespan of the current cohort has been reduced to 3.7 years which may be due to excessive fishing of juveniles as they attain attractable size at this stage as compared to other carps. However, all these growth parameters solely depend upon the largest fish size caught during the sampling period. Therefore, as the largest size caught during the period was comparatively smaller than earlier study, thereby, related parameters may have shown relatively lower range than those.

The probability of capture has shown that 50% of population measuring about 38 cm are prone to fishing activities. However, half of the mahseer population attain maturity at the length of 52 cm while majority of population attain maturity at 88 cm the (Nautiyal, 1985). This indicate that the most of the individuals of Mahseer get caught even before attaining the sexual maturity. The size-composition of present population also indicate distorted population structure as, the pre-reproductive size-groups measuring between 1 - 25 cm comprising about 78.4 % of the assessed stock. This population is dominated by size group measuring between 10 -15 cm, which alone accounted for approximately 24 %. However, larger size group mainly form the spawning biomass in the Nayar river as they migrate toward their breeding grounds solely for the purpose of spawning which explains their comparatively lower share in toral catch. Therefore, the over-exploitation of juveniles (sexually mature) and migratory brooders is essentially governing the phenomenon of “growth and recruitment overfishing”.

The depletion of juvenile and migratory cohorts, including those with potential for breeding, is a significant contributor to the reduction of the Mahseer population in the Indian subcontinent (Bhatt and Pandit, 2016). In addition to *T. putitora*, other species of *Tor*, such as *T. tor, T. duronensis*, and *T. tambroides*, are also at high risk of overexploitation (Ngguyen et al., 2007). In order to achieve optimal exploitation of the fish cohort, fishing mortality rates should not exceed 40% of natural mortality (i.e., F_opt_ = 0.4M), which would result in an optimal exploitation rate of 0.5/year. Here, the fishing mortality has surpassed the optimum level i.e., estimated as 0.42. The exploitation has also exceeded the optimum exploitation level as it was estimated as 0.67 which is far beyond the optimum. The earlier studies on the Gangetic stock have also recorded exploitation level beyond optimum which were estimated as 0.76 and 0.85 for 1980 and 1994 stock respectively (Nautiyal et al., 2008). This indicate that overexploitation has not receded in recent years even though several conservation activities are being employed under Mahseer conservation programmes.

In the context of management, biological reference points serve as a performance indicator for the fish stock being assessed. These points typically encompass a range of stock dynamics parameters, such as growth, recruitment, and mortality, with fishing mortality also taken into account, and subsequently translated into a consolidated index E0.1 is commonly favored over Emax as a biological reference point for the enduring management of fish stocks. To ensure optimal yield and biomass per recruit, the exploitation rate can be diminished by up to 0.278. The current exploitation also indicated the overexploitation as also reported in earlier stock reports of Mahseer (Nautiyal et al., 2008). As the Lc / L∞ ratio is less than 0.5, it signifies that juveniles constitute the major portion of the fishing cohort (Pauly and Munro, 1984), which is also reflected in the virtual population analysis. The graph of virtual population analysis further reveals the dominance of growth and recruitment fishing of Himalayan Mahseer as either peak was observed in juveniles and brooder group.

## Conclusion

The current classification of the Himalayan Mahseer as endangered by IUCN (2018) is a result of increasing habitat degradation owing to river management, hydro project construction, mining and quarrying activities, the discharge of pollutants, and rampant poaching. Consequently, the objective of the research was to comprehend the evolving growth parameters of the Himalayan Mahseer in the Nayar’s breeding and nursery grounds. The fundamental aim of managing fisheries is to ensure sustainable production over an extended period of time, primarily through regulatory and enhancement measures. The present investigation provides precise details regarding the status of the Himalayan Mahseer in the breeding and nursery grounds of the Western Himalayas. To enhance the present stock status and maintain the population of T. putitora in the foothill section of the Ganga River system (specifically the area between Rishikesh and Haridwar), it is recommended that fishing efforts on migratory brooders be reduced, and the mesh size be increased to prevent the exposure of sextually immature juveniles. The negligence of current stock characteristics of *T. putitora* could lead to collapse of foothill resident stock of Himalayan Mahseer distributed in Western Himalayan region in forthcoming future.

## Notes

### Competing Interest Statement

The authors have declared no competing interest.

### Summary of Updates

Some sort of data that were missed earlier are incorporated. The title of the manuscript was not going well with the manuscript so it has also been changed. Further, the section of results and discussion which were earlier incorporated together are placed separately.

